# Networks of Splice Factor Regulation by Unproductive Splicing Coupled With Nonsense Mediated mRNA Decay

**DOI:** 10.1101/2020.05.20.107375

**Authors:** Anna Desai, Zhiqiang Hu, Courtney E. French, James P. B. Lloyd, Steven E. Brenner

## Abstract

**Background:** Nonsense mediated mRNA decay (NMD) is an RNA surveillance pathway that degrades aberrant transcripts harboring premature termination codons. This pathway, in conjunction with alternative splicing, regulates gene expression post-transcriptionally. Nearly all serine and arginine-rich (SR) proteins and many heterogeneous nuclear ribonucleoproteins (hnRNPs) produce isoforms that can be degraded by the NMD pathway. Many splicing factors have been reported to be regulated via alternative splicing coupled to NMD. However, it is still uncharacterized that to what extent NMD contributes to the regulation of splicing factors.

**Results:** Here, we characterized a regulatory network of splicing factors through alternative splicing coupled to NMD. Based upon an extensive literature search, we first assembled a network that encompasses the current knowledge of splice factors repressing or activating the expression of other splicing factors through alternative splicing coupled to NMD. This regulatory network is limited, including just a handful of well-studied splicing factors. To gain a more global and less biased overview, we examined the splicing factor-mRNA interactions from public crosslinking-immunoprecipitation (CLIP)-seq data, which provides information about protein–RNA interactions. A network view of these interactions reveals extensive binding among splicing regulators. We also found that splicing factors bind more frequently to transcripts of other splicing factors than to other genes. In addition, many splicing factors are targets of NMD, and might be regulated via alternative splicing coupled to NMD, which is demonstrated by the significant overlap between the experimental network and eCLIP-network. We found that hierarchy of the splicing-factor interaction network differs from the hierarchy observed for transcription factors.

**Conclusion:** The extensive interaction between splicing factors and transcripts of other splicing factors suggests that the potential regulation via alternative splicing coupled with NMD is widespread. The splicing factor regulation is fundamentally different from that of transcription factors.

## Background

RNA splicing is regulated by RNA binding proteins, including serine and arginine-rich (SR) regulators, heterogeneous nuclear ribonucleoproteins (hnRNPs), and other splicing factors. Splicing factors affect virtually all human multi-exon genes that undergo alternative splicing to increases proteome diversity [1]. They dynamically regulate the transcriptome during growth and development in a cell-type-specific or tissue-specific manner [2, 3]. For example, PTBP2 is essential for neuronal maturation [4, 5], MBNL3 antagonizes muscle differentiation [6], and NOVA2 is a key regulator of angiogenesis [7]. Splicing factors maintain homeostasis but when unbalanced, can cause disease [8–11]. For example, defects in NOVA or TRDBP may result in severe neuronal pathogenesis [12–15].

Gene expression is regulated at multiple stages, from transcriptional initiation to post-translational modifications of the final protein product At the DNA level, transcription factors bind to cis-regulatory elements to modulate gene expression [16–19]. This regulation is often achieved in a combinatorial fashion through regulatory networks [20, 21]. These networks are often described as having a hierarchical architecture. Gerstein *et al.* employed a hierarchy height metric to measure the flow of information in a transcription factor network [22] and seperated transcription factors into three levels: a top-level “executive” that regulates many other factors, a middle level that co-regulates targets to mitigate information-flow bottlenecks, and a bottom-level “foreman” that is mostly regulated by other transcription factors. At the RNA level, gene regulation is controlled by splicing factors through binding to cis-regulatory RNA elements. Splicing factors have also been suggested to form hierarchical networks, with splicing factors positioned at the top of a splicing cascade forming “master splicing regulators” [23]. However, this definition of a master splicing regulator presupposes that the master regulator is obligatory for the proper differentiation or specification of a cell type. In addition, once a cell is committed to a lineage, this master regulator is required for maintaining homeostasis [23].

Regulated unproductive splicing and translation (RUST) is a mechanism that links alternative splicing with nonsense mediated mRNA decay (NMD) to regulate the abundance of mRNA transcripts [24, 25]. In this mode of regulation, unproductive alternative splicing introduces a premature termination codon into a transcript, which is consequently degraded by the NMD pathway. As a result, the productive transcript decreases in abundance, reducing the amount of the active gene product [26, 27]. Many transcripts undergo alternative splicing coupled to NMD in either a constitutive or regulated manner. RUST describes the latter and allows for gene expression to be fine-tuned depending on the cellular context. Many splicing factors are known to auto-regulate their own expression via RUST by binding their own pre-mRNA and promoting the unproductive transcript isoform [28–34]. This negative feedback loop regulates the expression of many splicing factors, ensuring that gene expression does not exceed desired levels. Splicing factors are also known to control the expression of other splicing factors through RUST. For example, PTBP1 (PTB) is expressed in non-neuronal tissues and in addition to regulating its own expression via RUST, normally acts to repress the expression of the productive transcript isoform of PTBP2 (nPTB). This ensures that the neuronal splicing program initiated by PTBP2 is not activated in non-neuronal cells. In neuronal cells, PTBP1 is repressed through the actions of a microRNA [5], leading to elevated levels of the productive isoform of PTBP2 and the consequent neuronal splicing events.

To estimate the extent of the splicing factor regulation via RUST, we collected experimentally validated cross-regulatory events through RUST between splicing factors (Table 1). Based on these reported RUST regulations, we reconstructed a splicing factor *regulatory* network. While this network encompasses hand-picked and well-studied splicing regulators, these were almost exclusively characterized through small-scale studies, resulting in a network that is largely underexplored. To enrich network content, we looked at studies mapping the interaction of splicing factors with mRNA, a necessary step in mRNA regulation. To study the extent of splicing factor-mRNA (SF-mRNA) interactions, we collected crosslinking-immunoprecipitation (CLIP) based studies [35], including CLIP-seq, iCLIP, PAR-CLIP, and eCLIP. Based on these collected splicing factor – splicing factor mRNA interactions, we recreated a splicing factor *interaction* network. All interactions in this network have the potential to be regulatory. However, there are caveats to our approach. The CLIP-seq experiment has low efficiency of the UV-mediated crosslinking, which ranges between 1% and 5%. Therefore, some interactions between a protein and its target RNA might not be captured, potentially leading to a less connected SF-mRNA interaction network. Also, this interaction network might contain non-physiological interactions since the collected CLIP studies were obtained from different cell types and were derived from two different species (humans and mice). Here, we demonstrated that an interaction network based on all collected CLIP studies contains 60% of all possible interactions, suggesting that regulation among splicing factors is potentially prevalent. When we focus only on the stringent eCLIP experiment [36], which uses input normalization controls and which is more uniform with respect to the experimental approach, the number of interactions declines to 30% of all possible interactions. Using only the eCLIP data, compared the hierarchy of splicing-factor interaction network to the hierarchy observed for transcription factors. In contrast to transcription factors, splicing factors do not form a hierarchical network, but instead display a nearly flat distribution across the hierarchy levels, with fewer factors acting at the extremes of the hierarchy.

**Table 1.** Literature based RUST regulations

| Source (Splicing Factor) | Transcript tested for NMD | Did the factor undergo CLIP-seq? | Regulation type | Is this auto-regulation ? | Organism | Reference | CLIP-confirmed |
| --- | --- | --- | --- | --- | --- | --- | --- |
| SRSF6 | SRSF1 | no | not observed | no | human | [28] | not tested |
| SRSF7 | SRSF2 | no | not observed | no | human | [29] | not tested |
| SRSF8 | SRSF2 | no | not observed | no | human | [29] | not tested |
| RBFOX3 | RBFOX2 | no | repressing | no | mouse | [37] | not tested |
| TRA2B | TRA2B | no | repressing | yes | human | [38] | not tested |
| PTBP1 | SRSF3 | yes | promoting | no | human | [39] | no |
| SRSF1 | SRSF2 | yes | not observed | no | human | [29] | yes |
| SRSF2 | SRSF1 | yes | not observed | no | human | [40] | yes |
| HNRNPL | SRSF3 | yes | repressing | no | human | [41] | no |
| PTBP1 | PTBP1 | yes | repressing | yes | human | [31] | no |
| PTBP1 | PTBP1 | yes | repressing | yes | human | [42] | no |
| FUS | SNRNP70 | yes | promoting | no | mouse | [43] | yes |
| PTBP2 | SRSF3 | yes | promoting | no | human | [39] | yes |
| RBFOX2 | SNRNP70 | yes | promoting | no | mouse | [32] | yes |
| RBFOX2 | SRSF7 | yes | promoting | no | mouse | [32] | yes |
| RBFOX2 | PTBP2 | yes | promoting | no | mouse | [32] | yes |
| RBFOX2 | TIA1 | yes | promoting | no | mouse | [32] | yes |
| SRSF1 | SRSF3 | yes | promoting | no | mouse | [44] | yes |
| SRSF4 | SRSF1 | yes | not observed | no | human | [28] | no |
| SRSF4 | SRSF2 | yes | not observed | no | mouse | [45] | no |
| SRSF4 | SRSF3 | yes | not observed | no | mouse | [45] | no |
| SRSF4 | SRSF5 | yes | not observed | no | mouse | [45] | no |
| SRSF4 | SRSF7 | yes | not observed | no | mouse | [45] | no |
| SRSF9 | SRSF1 | yes | not observed | no | human | [28] | no |
| ELAVL1 | PTBP2 | yes | repressing | no | human | [46] | yes |
| HNRNPA1 | HNRNPA2B1 | yes | repressing | no | human | [47] | yes |
| HNRNPL | HNRPLL | yes | repressing | no | human | [30] | yes |
| KHDRBS1 | SRSF1 | yes | repressing | no | human | [48] | yes |
| PTBP1 | PTBP2 | yes | repressing | no | human | [42] | yes |
| RBFOX2 | TRA2A | yes | repressing | no | mouse | [32] | yes |
| RBFOX2 | SF1 | yes | repressing | no | mouse | [32] | yes |
| SRSF3 | SRSF7 | yes | repressing | no | mouse | [45] | yes |
| SRSF3 | SRSF2 | yes | repressing | no | mouse | [45] | yes |
| SRSF3 | SRSF5 | yes | repressing | no | mouse | [45] | yes |
| RBM10 | RBM5 | yes | repressing | no | human | [40] | yes |
| TIA1 | TIA1 | yes | repressing | yes | mouse | [32] | yes |
| HNRNPA2B1 | HNRNPA2B1 | yes | repressing | yes | human | [49] | yes |
| HNRNPL | HNRNPL | yes | repressing | yes | human | [30] | yes |
| PTBP2 | PTBP2 | yes | repressing | yes | mouse | [32] | yes |
| SRSF1 | SRSF1 | yes | repressing | yes | human | [40] | yes |
| SRSF2 | SRSF2 | yes | repressing | yes | human | [29] | yes |
| SRSF3 | SRSF3 | yes | repressing | yes | mouse | [45] | yes |
| SRSF3 | SRSF3 | yes | repressing | yes | mouse | [44] | yes |
| RBM10 | RBM10 | yes | repressing | yes | human | [40] | yes |
| FUS | FUS | yes | repressing | yes | human | [50] | yes |

## Results

### Literature-based RUST network of splicing factors

The expression of many splicing factors is controlled via RUST. We collected published regulations between splicing factors through RUST (as described in Methods) (Table 1) [28–32, 37–50]. We identified 19 publications reporting 19 splicing factors as targets of RUST in human and murine cells. These included five SR factors (SRSF1, SRSF2, SRSF3, SRSF5, and SRSF7), three hnRNPs (hnRNPA2B1, hnRNPL, and hnRNPLL), and eleven other splicing regulators (RBFOX2, RBFOX3, FUS, SNRNP70, TRA2A, TRA2B, TIA1, PTBP1, PTBP2, RBM10, and RBM5) (Table 1). Eleven splicing factors auto-regulate their own expression through RUST: SRSF1, SRSF2, SRSF3, PTBP1, PTBP2, hnRNPL, hnRNPA2B1, TIA1, TRA2B, FUS, and RBM10 (Figure 1). All auto-regulatory RUST events are repressive. In the repressive mode of RUST, a splicing factor binds to a transcript and promotes alternative splicing changes that favor production of the unproductive transcript. For example, PTBP1 binds to its own pre-mRNA and promotes an alternative splicing change that leads to the exclusion of exon 11, creating a transcript that is consequently degraded by the NMD pathway [31]. Fourteen targets of RUST are cross-regulated by other splicing factors: SRSF1, SRSF2, SRSF3, SRSF5, SRFS7, hnRNPL, hnRNPA2B1, PTBP2, RBM5, RBFOX2, TIA1, TRA2A, SF1, and SNRNP70 (Figure 1 and Table 1). Twelve of these targets are regulated by a single splicing factor. For example, RBFOX is regulated by RBFOX3. A caveat for many of these studies is that for the RUST regulation to be observed, the regulatory factor (in this example, RBFOX3) has to be overexpressed many times over its physiological levels with concurrent inhibition of the NMD pathway [37]. Just two splicing factor targets of RUST, PTBP2 and SRSF3, have been shown to be regulated by more than one splicing factor. Both splicing factors are promoted and repressed by other splicing factors: RBFOX2 advances expression of productive PTBP2, whereas, PTBP1 advances expression of unproductive PTBP2. SRSF1 advances expression of productive SRSF3, whereas, hnRNPL advances expression of unproductive SRSF3 (Figure 1). In the promoting mode of RUST, a splicing factor binds a transcript and advances alternative splicing changes that favor splicing of the productive transcript. Conversely, in the repressive mode of RUST, a splicing factor binds a transcript and advances alternative splicing changes that favor splicing of the unproductive transcript. For example, hnRNPL represses the expression of SRSF3 [41], whereas SRSF1, PTBP1, and PTBP2 promote the expression of SRSF3 [39, 44]. In another example, the PTBP2 productive transcript is promoted by RBFOX2 and repressed by PTBP1 and ELAVL1 [32, 33, 46]. Overall, out of the 21 cross-regulatory events reported, 8 are promoting and 13 are repressive (Figure 1 and Table 1).

**Figure 1.**
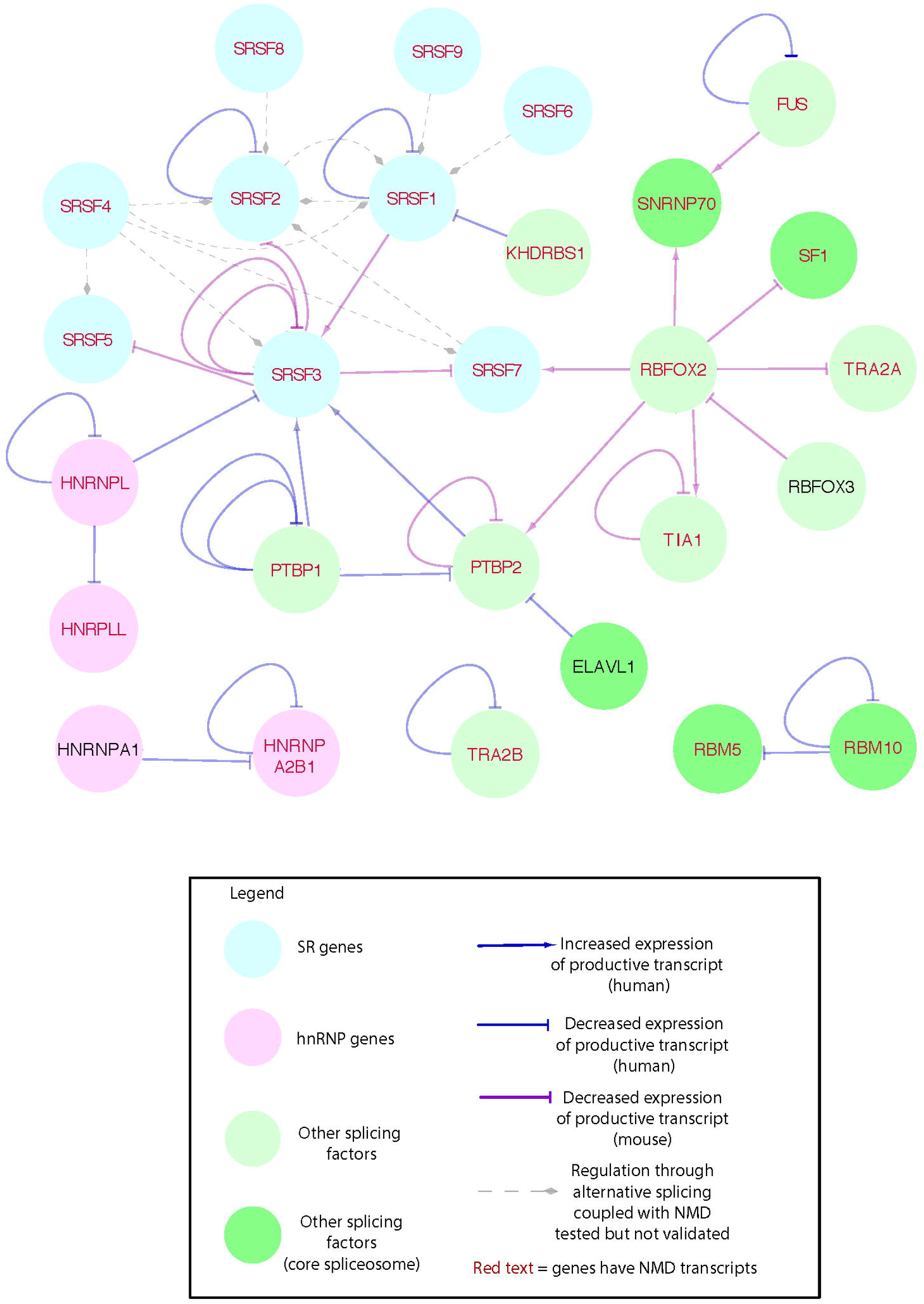
Splicing factor *regulatory* network. Each node represents a splicing factor and its transcript. Node color represents a family of splicing regulators as described in the legend. Red font represents a gene that has been shown to express an NMD targeted transcript. Nodes colored with a darker shade of green represent splicing factors that associate with the spliceosome. Edges represent RUST *regulations*. A node at the source of an edge represents a splicing factor, and a node at the target of the edge represents a target of RUST. Solid line edges represent the presence of RUST. Dashed line edges represent RUST regulation that was tested but not observed. Edges ending with a ⊥ represent gene repression via RUST. For example, PTBP1 (PTB) represses the expression of the productive transcript isoform of PTBP2 (nPTB); this regulation is depicted as an edge that originates at the PTBP1 node and ends with a ⊥ in the PTBP2 node. Edges that end with an ↟ represent gene promotion via RUST. Blue edges stand for RUST observed in humans, and purple edges stand for RUST observed in mice. Each edge represents a single experiment as listed in Table 1. This network consists of 27 nodes and 45 edges.

Some tested regulations have revealed no RUST. For example, overexpression of SRSF4 does not alter splicing patterns of SRSF2 [45]. Overall, RUST was observed in 76% (95% confidence interval, 61%~86%) of the tested cases (34/45), indicating that regulations via RUST are extensive, at least among splicing factors. It should be noted that this rate could be overestimated, due to a bias towards publishing positive results in the form of observed regulations, and a likely bias towards reporting factors that have important roles in splicing and biological processes.

### Extensive RUST revealed by a splicing factor interaction network

Almost all the above studies identified RUST on a small scale. In a typical study, a selected splicing factor is overexpressed or knocked down; and changes in alternative splicing, on a limited number of transcripts, are observed via semi-quantitative PCR, usually in an NMD-inhibited background. Only one study assessed RUST regulation on a wider scale. RBFOX2 has been shown to activate TIA1, PTBP2, SRSF7, and SNRNP70 and repress SF1 and TRA2A via RUST [32]. Here, RBFOX2 controls many splicing factors, but this could represent a bias since it is the only splicing factor whose regulation was tested on a genome-wide scale.

To examine if there is a potential for the regulatory network to be more extensive, we explored the binding of splicing factors and their target RNA, which is the prerequisite that a regulation occurs. To better understand the potential of regulation between these factors, we examined the transcriptome-wide binding patterns of splicing factors by exploring ENCODE eCLIP experiments. In these experiments, 161 RNA binding proteins were assayed in the same laboratory, using the same protocol, employing two cell lines: either HepG2 or K562 [33, 36, 51]. We took the stringent set of interactions generated by eCLIP, as summarized in Methods and as described in Van Nostrand *et al*. [36], to reduce the number of false positive interactions and to keep the results consistent. Here, we focused on eCLIP, rather than on all available CLIP experiments, to reconstruct splicing factor-mRNA interactions that are most likely physiological since they are observed in the same cell line (K562 or HepG2). Also, the eCLIP results obtained from assaying multiple splicing factors likely can be compared directly, since all individual experiments follow the same protocol. Finally, all eCLIP experiments employ normalization steps, paired size-matched input (SMInput) and IgG-only controls [36], which increased our confidence. We identified genome-wide RNA binding sites of a splicing factor using stringent parameters (p-value <0.05 and >8 fold enrichment, see Methods for details) and then extracted the binding sites of mRNAs from the splicing factors. The stringency of eCLIP is also its limitation since in some cases it severely reduces the number of SF-mRNA interactions for certain splicing factors. For example, eCLIP of hnRNPA1 in HepG2 cells provides only 40 splicing factor-mRNA interactions (Table S1), whereas CLIP-seq of hnRNPA1 in HEK293T cells identifies 2,043 SF-mRNA interactions (Table S2) [52].

We selected 21 and 22 RNA binding proteins that have been experimentally shown to alter alternative splicing and that underwent eCLIP in HepG2 and K562 cells, respectively (a total of 27 unique splicing factors) (Figure 2A-B and Table S1). Among these selected RNA binding proteins, we included factors that are core splicing factors but which also have been shown to alter alternative splicing. These include U2AF1, U2AF2, SF3A3, and SF3B4 [53–57]. Out of these 27 unique eCLIP assayed factors, 16 splicing regulators were also studied in both cell lines. The collection of confident eCLIP binding peaks (interactions between a splicing factor and any part of the bound gene) were converted into network edges and illustrated in two separate networks: one for K562 and one for HepG2 cells. Overall, 31.6% (153 out of 484) of possible interactions were present in the K562 eCLIP network and 30.6% (135 out of 441) of possible interactions were present in the HepG2 eCLIP network (Figure 2A-B). On average, a splicing factor in the eCLIP K562 network had 13.9 edges, with a wide range from 2 for hnRNPK to 29 for U2AF2, whereas a splicing factor in the eCLIP HepG2 network had 12.9 edges, ranging from 3 for SRSF9 to 23 for DDX3X. A splicing factor in the eCLIP K562 network had an average of 11.5 neighbors out of 21 possible, whereas a splicing factor in the eCLIP HepG2 network had an average of 10.9 neighbors out of 20 possible. There were seven self-loops in the K562 network and six self-loops in the HepG2 network. The presence of these self-loops implies that a third of splicing regulators might undergo auto-regulation. The high proportion of interactions suggest that splicing factors form a highly connected network of interactions, and this might suggest that cross-regulation, including that through RUST, is prevalent.

**Figure 2.**
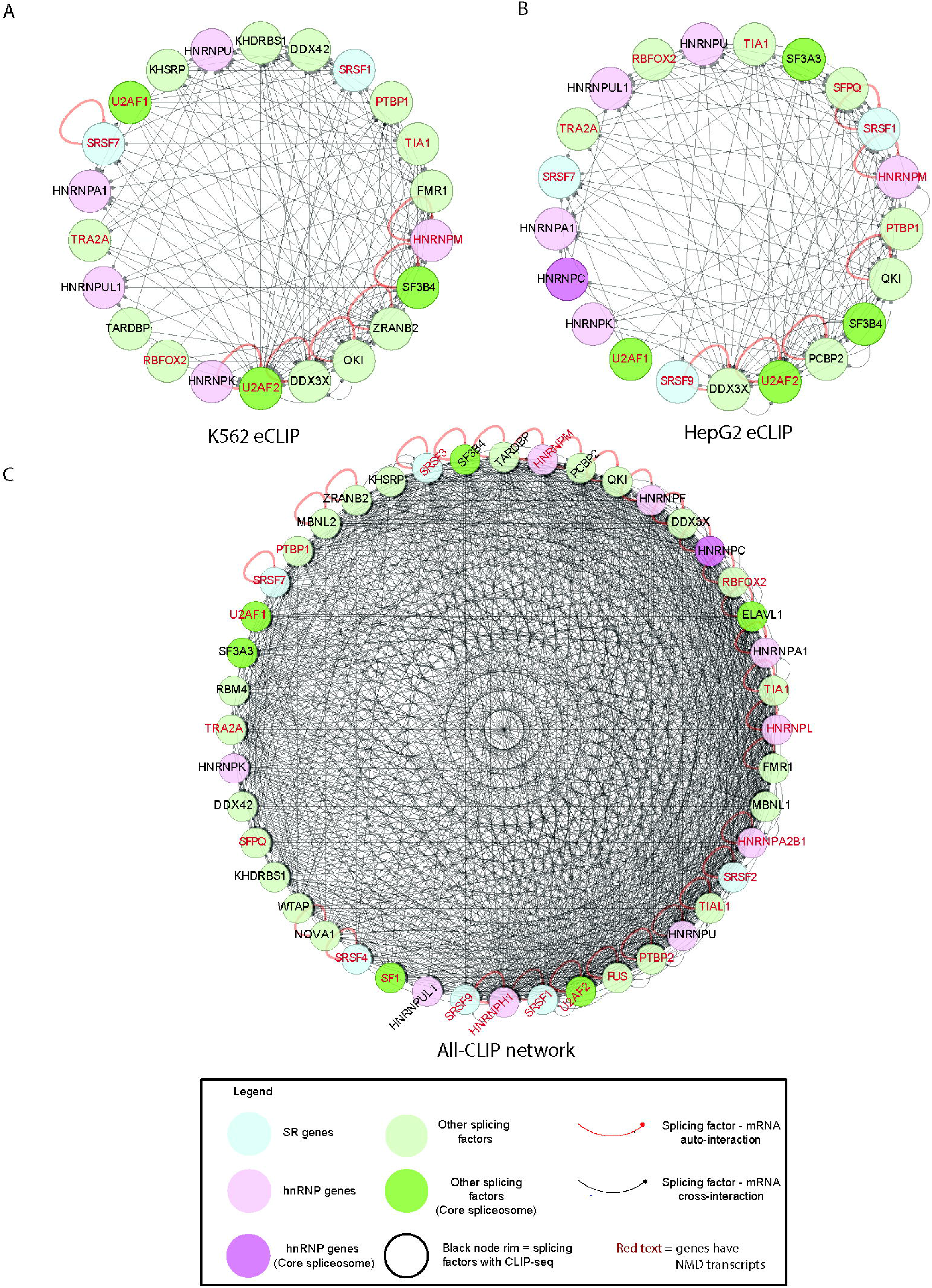
Splicing factor-mRNA interaction network. **A–B.** eCLIP detected network of interactions between splicing factors and their transcripts in K562 (**A**) and HepG2 (**B**) cells. Description of nodes is the same as in Figure 1. Edges represent splicing factor-mRNA interactions. A node at the source of the edge represents a splicing factor that underwent CLIP, and a node at the target of the edge, which ends with a dot, represents an mRNA transcript with which a splicing factor interacts. Multiple CLIP binding peaks located in a given transcript are portrayed as a single edge between a splicing factor and its mRNA target. Self-edges are colored red. The K562 based network contains 22 nodes and 152 interactions; the HepG2 based network contains 21 nodes and 135 interactions. The clustering coefficient is 0.46 for the K562 network and 0.41 for the HepG2 network. **C**. The all-CLIP based network contains 44 nodes and 1,153 edges; the clustering coefficient is 0.67. All networks are arranged in a degree sorted circle layout.

There was a large variation in the number of eCLIP-binding peaks identified for each factor (Table S1). For example, a genome-wide eCLIP of hnRNPA1 identified only 46 interactions, corresponding to transcripts of 39 genes, of which one (SFPQ) is a splicing factor from our list of 100 alternative splicing factors of interest in K562 cells. In HepG2 cells, an eCLIP of hnRNPA1 yielded 40 interactions, corresponding to transcripts of 19 genes, out of which only 2 are splicing factors (SF3B1 and SRSF6) from our list of 100 alternative splicing factors. Here, the eCLIP did not capture the established self-interaction of hnRNAPA1 [52, 58–60]. The highest number of eCLIP binding peaks was identified for DDX3X: 8,422 interactions in K562 cells, corresponding to 4,379 genes, of which 48 are splicing factors; and 9,644 interactions in HepG2 cells, corresponding to 4,183 genes, of which 38 are splicing factors. For many assayed splicing factors, eCLIP resulted in a similar number of binding peaks in the two different cell lines; this is true for hnRNAPA1, DDX3X, hnRNPM, and PTBP1 (Table S1). There were several factors for which eCLIP identified nearly twice as many factor-mRNA interactions in the HepG2 cells than in the K562 cells. These factors included: hnRNPK, hnRNPU, hnRNPUL1, QKI, RBFOX2, and U2AF2 (Table S1). Conversely, splicing factors SF3B4, TIA1, TRA2A, and U2AF1 exhibited significantly fewer SF-RNA interactions in the HepG2 cells than in the K562 cells (Table S1). These observations indicate that many splicing factors have cell line-specific targets, and consequently might regulate different splicing factors in different cell lines.

In addition to eCLIP, we gathered an additional 46 CLIP experiments, representing 34 splicing factors, to assess the extent of interactions between splicing factors and their transcripts among a more diverse sets of experiments. Here, we collected interactions that were deemed confident in their original publications. These additional studies included 27 CLIP-seq, 11 iCLIP, and 8 PAR-CLIP experiments (for a detailed list, see Table S2). Thirty-one were performed in human cells or tissues, and 15 were conducted in murine cells or tissues (Table S2). These splicing factor-mRNA interactions were merged with the eCLIP data. The resulting splicing factor-mRNA interaction network (Figure 2C), named all-CLIP network, contains 44 splicing factors (nodes) and 1,153 interactions (edges). The edges comprise nearly 60% of all possible network interactions (1,153 present out of 1,936 possible). Twenty-nine of these interactions are self-interactions which might be auto-regulatory. In this network, on average, each splicing factor has 35.7 neighbors, and the clustering coefficient of this network is 0.67. This highly connected network is twice as dense as the eCLIP derived network, further hinting that RUST has the potential to be prevalent. This all-CLIP network comes with many caveats, such as a large variation in identified CLIP clusters even when the same splicing factor is crosslinked to RNA (see Table S2 for details) and diverse data analyses among studies. In addition, these studies were performed using different biological materials yielding a variable number of interactions. For example, hnRNPA1 underwent iCLIP in HeLa cells, which identified 40,670 interactions [60], whereas CLIP-seq of hnRNPA1 in HEK293T cells identified 2,043 interactions [52]. Hence, by combining experiments from different cell lines and tissues, as well as combining murine and human interactions, we likely introduced many false positive splicing factor–mRNA interactions. Despite the abovementioned caveats, this network indicates a sizeable potential for cross-regulation between splicing factors and their transcripts.

To further analyze the potential of the RUST interaction network among an extended list of splicing regulators, we added 56 alternative splicing factors to the all-CLIP network. These additional splicing factors have not undergone CLIP but have been shown to affect alternative splicing (see Methods). This expanded network reveals that the transcripts of all 100 splicing factors are bound by another splicing regulator. The numbers of these incoming interactions vary, ranging from 4 for CELF3, to 35 for hnRNPH1 (Figure 3). In this network, many splicing factors interact with transcripts of over 80 splicing regulators. These factors include: SRSF2 (82), TIA1 (82), hnRNPH1 (83), SRSF1 (84), U2AF2 (85), TIAL1 (86), RBFOX2 (86), PTBP2 (88), hnRNPC (89), FMR1 (89), and hnRNPL (92). Only a handful of splicing regulators bind 10 or fewer splicing regulator transcripts. These factors include: SFPQ (1), hnRNPUL1 (2), SF1 (5), DDX42 (6), SRSF9 (8), and KHDRSB1 (9). This extended all-CLIP network contains 2,180 interactions. Here, on average, a splicing factor has 36 neighbors, and many splicing factors express NMD-targeted transcripts (40, red font nodes in Figure 3) [61]. This highly connected network of 100 splicing factors indicates that many more splicing factors might be regulated through RUST.

**Figure 3.**
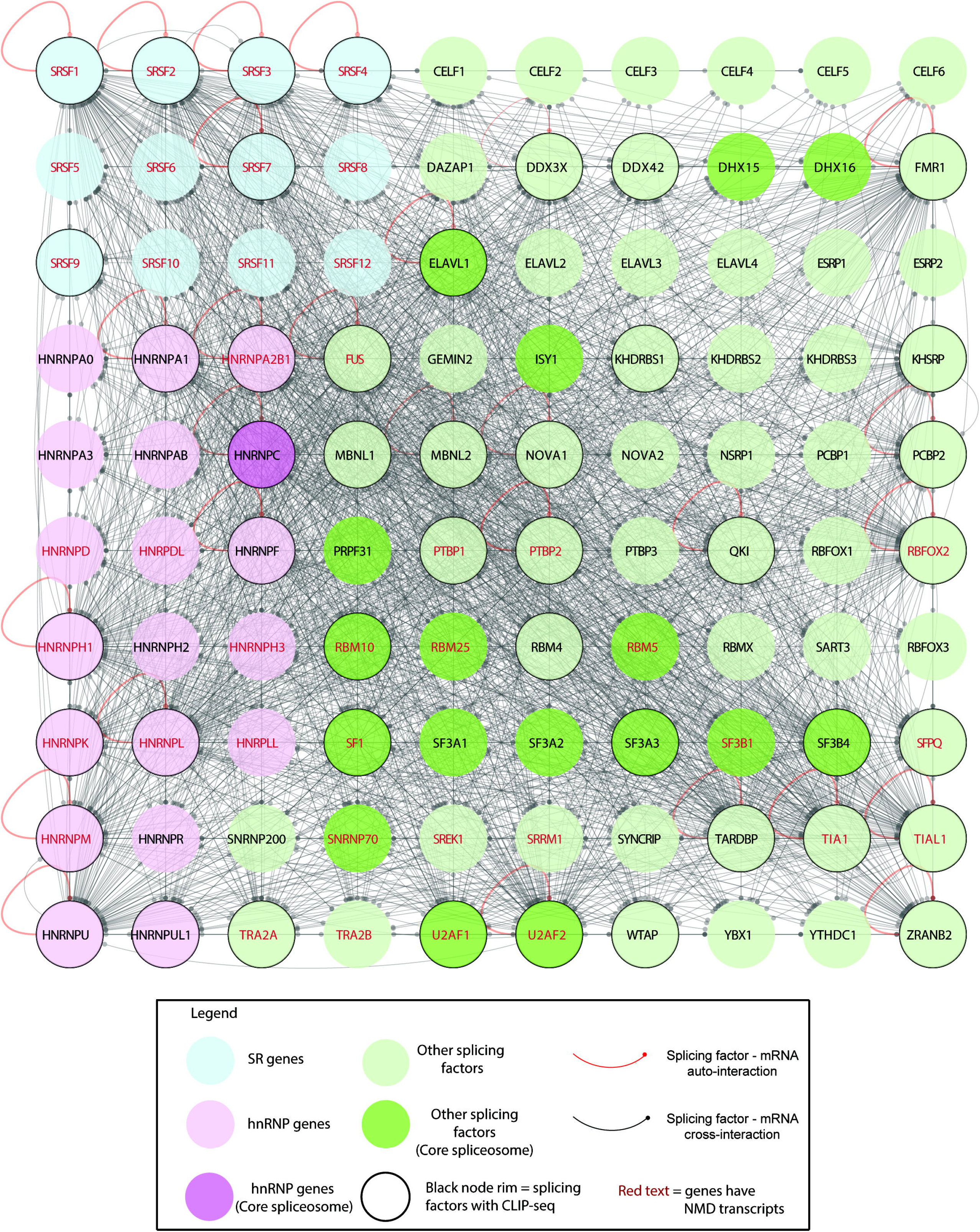
Pictorial representation of agreement or lack of it between the RUST regulatory network and the All-CLIP network. Nodes represent splicing factors (as described in Figure 1). Solid edges are marked in green when there are supporting CLIP interactions. Dashed edges are market in green when CLIP interactions are absent. Red edges identify interactions that are in conflict with CLIP. For example, hnRNPL represses SRSF3, but interaction between hnRNPL and SRSF3 have not been confirmed. Orange edges identify regulations that have been neither supported or refuted by CLIP due to lack of CLIP experiments. For example, RBFOX3 did not undergo CLIP.

Without more experimental data, it is difficult to determine what fraction of splicing factor-mRNA interactions might represent RUST. However, one can compare the experimentally proven RUST network to the available CLIP binding peaks to further infer if CLIP interactions could be used to predict regulation (Figure 3 and Table 2). Overall, there is a statistically significant agreement between the CLIP interactions and RUST (p-value of 0.0004; two-tailed Fisher test). RUST was observed in 34 experiments and was confirmed by 29 interactions (85% agreement) (Table 2, green solid edges in Figure 3). Twenty-eight interactions were confirmed by CLIP and one (TRA2B) was confirmed by Stoilov *et al.* [38]. More important, when RUST was not observed (11 instances), 7 CLIP interactions were absent (64% agreement, green dashed edges in Figure 3) with only 2 false positives in which CLIP detected interaction but for which experimental RUST was not confirmed (Table 2, red dashed edges in Figure 3). This overlap hints that approximately 60% of CLIP based interactions in our network (Figure 3) might present potential regulations via RUST.

**Table 2.** Overlap of RUST and CLIP

| **CLIP** | **RUST observed** | **RUST not observed** |
| --- | --- | --- |
| **yes** | **29** (green solid edges in Fig. 3) | **2** (red dashed edges in Fig. 3) |
| **no** | **4** (red solid edges in Fig. 3) | **7** (dashed green edges in Fig. 3) |
| **not tested** | **1** (orange solid edge in Fig. 3) | **2** (orange dashed edges in Fig. 3) |
| **total** | **34** | **11** |

### Different hierarchy between the splicing factor regulatory work and the transcription factor regulatory network

Transcription factors form hierarchical networks [22, 62, 63]. To determine if splicing factors form a similar hierarchical network, we used the same approach as Gerstein *et al.* [22]. The hierarchy height of each transcription factor was determined by the incoming and outgoing edges; all transcription factors were classified into three layers based on their hierarchy height (see Methods for detail). The top layer is for “executive” regulators and factors with a value in the top third of possible hierarchy height values. The middle layer is for factors with values from the center of this range, and “foremen” are from the lower third of this range. We applied the same hierarchy height metric to the ENCODE eCLIP data (Figure 4 and Supplementary Figure 1). Transcription factors were highly peaked at 1 and −1 values (Figure 4A and Supplementary Figure 1A) and distinctly grouped into three layers (see the three hierarchy height peaks on Figure 4C and Supplementary Figure 1C). By contrast, splicing factors were nearly uniformly abundant across the entire hierarchy height axis (Figure 4B, D and Supplementary Figure 1B, D). The hierarchies were robust in different cell lines for both the transcription factors (Figure 4C and Supplementary Figure 1C, p-value = 0.88, two-sided chi-square test) and the splicing factors (Figure 4D and Supplementary Figure 1D, p-value=0.72, two-sided chi-square test). There was significant difference between the hierarchies of transcription and splicing factors (Figure 4C–D, p-value = 0.031, two-sided chi-square test). Interestingly, nearly all splicing factors with auto-loops gathered in the middle layer (6/7 in K562 as shown in Figure 4 and 5/5 in HepG2 as shown in Figure S1). The only exception was SF3B4, which is an executive factor but had a lower score among the splicing factor executive layer. Moreover, in the K562 eCLIP based network, two splicing factors, SRSF1 and PTBP1, are middle level regulators, and do not have evidence for auto-regulation within the eCLIP data set. However, they have been demonstrated to autoregulate in other studies [28, 31].

**Figure 4.**
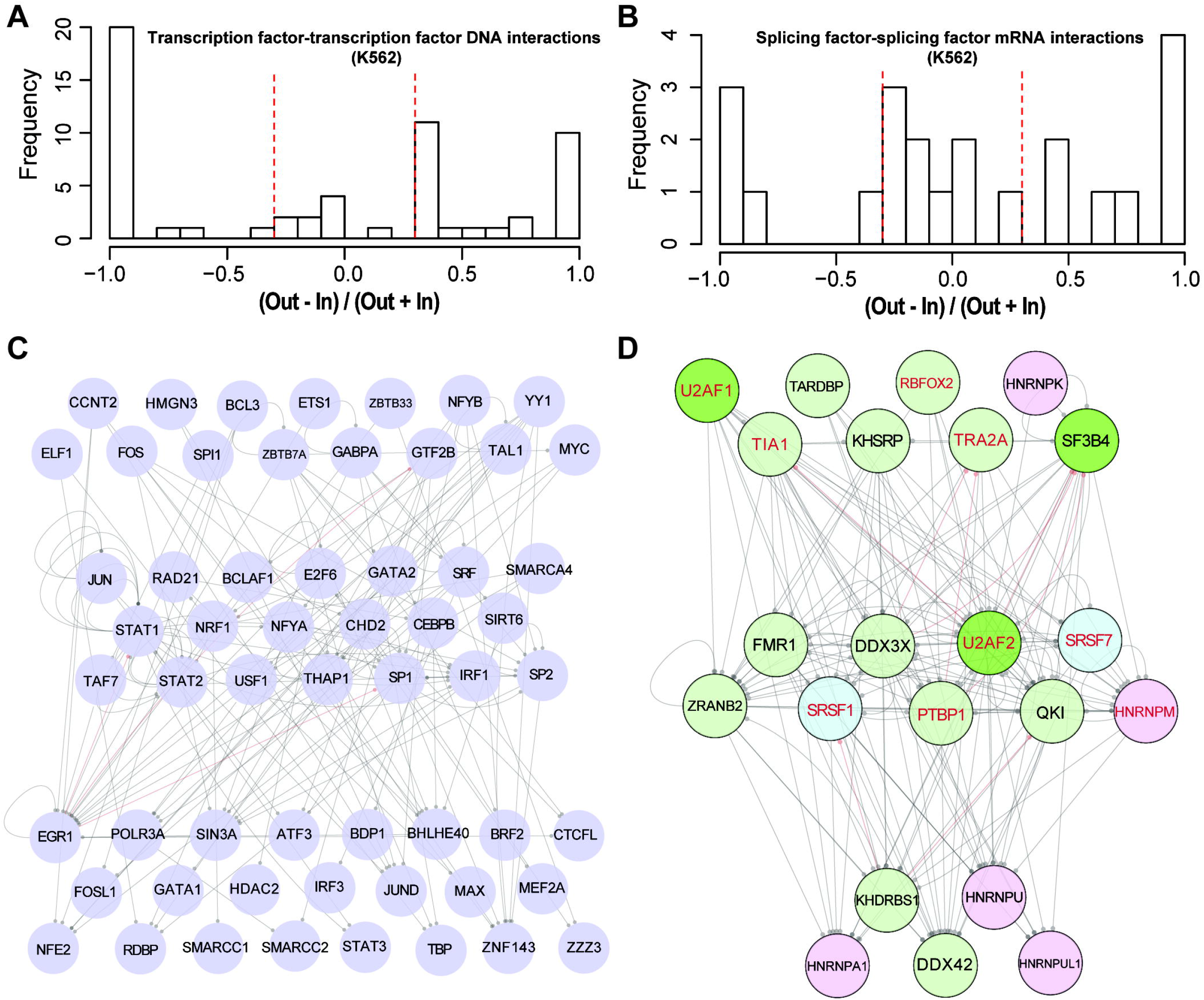
Comparison of hierarchies of the transcription factor–transcription factor DNA interaction network and splicing factor–splicing factor mRNA interaction network in the K562 cell line. **A–B.** The distributions of node hierarchies defined as h = (Out – In)/(Out + In), where “Out” represents the out-degree and “In” represents the in-degree, in the transcription factor–transcription factor DNA interaction network (**A**) and splicing factor–splicing factor mRNA interaction network (**B**). **C–D**. The transcription factor–transcription factor DNA interaction network (**C**) and the splicing factor–splicing factor mRNA interaction network (**D**) shown in a hierarchy. Nodes with h >1/3 and h < −1/3 are on the top and bottom level, respectively, with the other nodes located in the middle. The grey lines show top-down interactions while red lines show bottom-up interactions. The colors are the same as in Figure 2.

To identify master regulators at the top of the hierarchy with the ability to directly regulate other splicing factors, we looked for splicing factors that bound to more transcripts of other splicing factors than to transcripts of non-splicing factor other genes. We compared the frequency of splicing factors binding to transcripts of all genes versus those binding to transcripts of other splicing factors (Figure 5 and Table S3). This comparison revealed that there are no clear outliers. On average, a splicing factor binds transcripts of other splicing regulators five times more frequently than transcripts of all other expressed genes (Figure 5 and Table S3). While all splicing factors follow this trend, there is a wide range. For example, SRSF7 and KHDRBS1 bind transcripts of splicing factor genes 12 times more frequently than transcripts of all other genes in the K562 cells, whereas, hnRNPU binds transcripts of splicing factor genes 11 times more frequently than transcripts of all other genes in the HepG2 cells. Also, hnRNPA1 exhibits preferential binding to transcripts of other splicing factors than to transcripts of all other genes (17 times more) in the HepG2 cells; however, this result is based on a small overall number of eCLIP clusters identified for this factor and might be an aberration. At the other end of the spectrum, factors such as PTBP1 and DDX3X show only slight preference for binding transcripts of splicing factor genes than transcripts of all other genes. For example, PTBP1’s preference reaches 1.3 in both cell lines (Table S3). Interestingly, a combined plot for all regulators that underwent eCLIP (43 studies) reveals that there is a continuum in the preferential splicing factor binding to transcripts of other splicing factors, with no clear master regulator.

**Figure 5.**
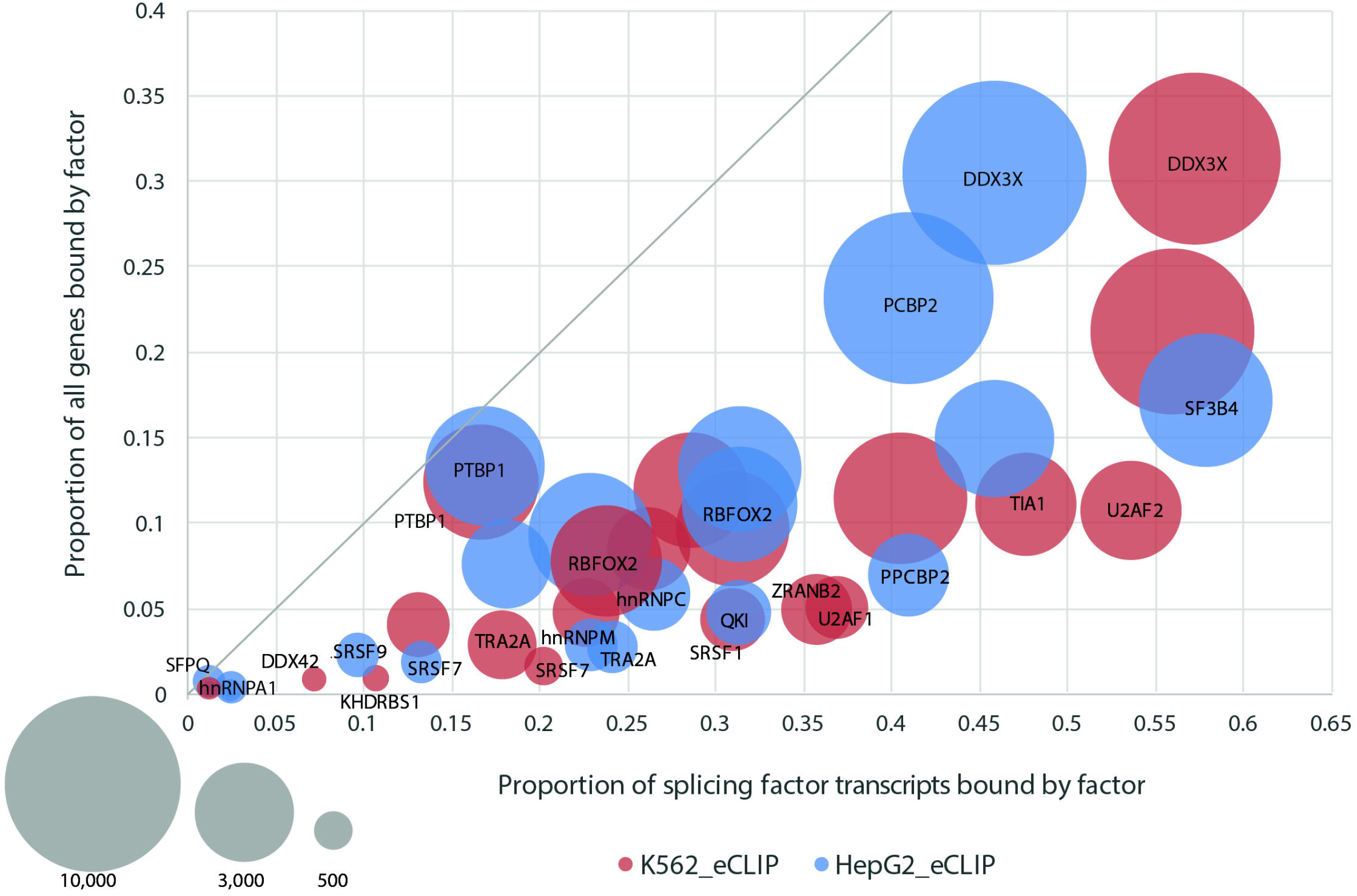
All evaluated splicing factors bind transcripts of other splicing factors more prevalently than transcripts of all other genes. Plotted are ratios of the number of splicing factors a protein binds over the number of all expressed splicing factors in a given cell line (100 selected splicing factors) versus the ratio of the number of transcripts a splicing factor binds over transcripts all expressed genes in a given cell line with an FPKM value above 1. The data were obtained from eCLIP experiments. The size of each circle represents the total number of CLIP clusters obtained by eCLIP. Names of the selected factors are noted on the plot. For more details see Table 4.

## Discussion

While regulation of gene expression has been studied extensively at the transcription factor level, regulation of gene expression by splicing factors has been studied to a lesser extent. Individual splicing factors have crucial influence over cell fate and development [64, 65]. A survey of the literature reveals that expression of several splicing factor genes is regulated via RUST (Table 1). This network seems sparse, probably because it is biased towards well studied splicing regulators. To regulate alternative splicing, splicing factors are required to interact with pre-mRNA of their target. Hence, to gain a global view of such regulations, we gathered publicly available splicing factor–mRNA interactions and organized them into a splicing factor–mRNA interaction network for 100 selected splicing regulators (Figure 3). This network is highly connected, suggesting that the potential for RUST is high.

Master regulators have been suggested to exist among splicing factors, however, the definition of a master regulator varies considerably. One such definition states that a splicing factor master regulator is a protein that maintains a specific cell lineage [23]. Under this definition, SRSF6 is a master regulator of eye development in *Drosophila* [66]. The entire SR protein family has also been described as master splicing regulators, based on the finding that their disruption leads to developmental defects and disease [67]. In another definition, a master regulator is a gene that occupies the very top of a regulatory hierarchy [68]. Here, we explore the hierarchy of splicing factors using the “hierarchy height” metric, as used by Gerstein *et al.*, and compare it to a hierarchy among transcription factors [22]. We show that the relationship among splicing factors differs from that of transcription factors. Transcription factors seem to fall naturally into three layers. In contrast, our limited data show that splicing factors are less easily split into three distinct groups. In addition, transcription factors with self-interactions can be spotted in all three layers, whereas splicing factors with self-interactions nearly exclusively group in the middle layer.

In the search for a unified regulatory splicing factor network, we found that splicing factors bind transcripts of other splicing factors, on average, five times as frequently as transcripts of other genes (Figure 5 and Table S3). This finding is in line with published studies that describe abundant splicing factor binding to transcripts of other splicing factors, as reviewed in [69] and [70].

Many factors could have affected our study. While CLIP is a powerful method that provides experimental interactions of an RNA binding protein with RNA, CLIP experiments suffer from low crosslinking efficiency [71] and might result in false negative results. Also, non-specific antibody–mRNA interactions might yield false positive results. We compared the overlap of the CLIP interactions to the RUST regulatory network, and nonetheless, there was a high overlap, with modest frequency of false positive and false negatives (Table S3). Therefore, despite limitations, we found that CLIP-seq seems to be a reliable method to predict regulations among splicing factors.

## Conclusion

We generated a literature-based network of splicing factor regulation through alternative splicing coupled to NMD. Using CLIP-seq and related studies, we found that this type of regulation could be much more prevalent than previously thought. Within these networks, the information flow among splicing factors differs from that described for transcription factor regulatory networks. Future RNA-seq experiments performed in biological systems with inhibited NMD pathway are necessary to confirm the full extent and significance of RUST in splicing factor regulation.

## Methods

### Splicing factor regulatory network

Through an extensive literature search through PubMed using “NMD regulation” and “splicing factor” as query terms, we collected 45 publications that described experimental results (Table 1) that tested overexpression or knockdown of a splicing factor resulting in alternative splicing changes of the ratio of productive to unproductive transcripts of self or other splicing factors.

The unproductive transcripts have been shown experimentally to be targets of the NMD pathway [28, 29, 31–33, 37–41, 43, 44, 47, 48, 60, 63, 72].

### Selection of 100 alternative splicing factors

The selection of 100 alternative splicing regulators was derived from the National Center for Biotechnology Information (NCBI) database of genes (http://www.ncbi.nlm.nih.gov/gene/) and the GeneCards^®^ database (http://www.genecards.org). We also performed an NCBI PubMed (www.ncbi.nlm.nih.gov) abstracts search for alternative splicing regulators. As a query, we paired an “alternative splicing” term with a list of RNA binding proteins included in the GO:0003723 Gene Ontology Consortium database (geneontology.org). The full list of splicing factors selected is presented in the form of a splicing factor–RNA interaction network (Figure 3).

### Splicing factor interaction network

We performed a literature search in PubMed and a Google search for CLIP-seq and related studies. We came across and collected 89 CLIP studies that used a splicing factor as the crosslinked protein; 74 were performed in human cells or tissues and 15 were conducted using murine biological material. The data (splicing factor–mRNA interactions) were obtained from accession numbers included in the above-mentioned references or from supplemental tables. The results are summarized in Table S1. The splicing factor–RNA interaction binding sites identified in each study were used to construct the networks presented in Figure 2 and Figure 3. When original studies provided only genomic coordinates of CLIP-clusters, we used appropriate Ensembl GTF annotation files (hg19, GRCh38, mm9) to add corresponding gene names. The collection of 89 CLIP studies includes 43 eCLIP studies that were downloaded from the ENCODE website (www.encodeproject.org). For eCLIP data, we downloaded two biological replicates (narrowPeak in BED format) for each splicing factor of interest (for accession codes, see Table S2). The two replicates were merged. Overlapping eCLIP clusters were collapsed using Yeo Lab Perl script (https://github.com/YeoLab/gscripts/tree/master/perl_scripts/compress_l2foldenrpeak.fi.pl). The clusters were filtered for significant clusters that are defined as splicing factor-mRNA binding sites that are identified as significant (*P* < 0.05) by CLIPper and are eightfold enriched above SMInput [46].

Cytoscape 3.4.0 [73] was used to visualize and analyze networks. We input tables with all interactions (individual tables were created for individual networks) into Cytoscape. We organized nodes using a custom setting or degree sorted circle. We also defined a custom style (for nodes) to visualize networks.

### Splicing and transcription factors hierarchical networks

The Transcription Factor–gene networks for GM12878 and K562 cell lines were downloaded from Gerstein *et al.’s* work [22] (http://encodenets.gersteinlab.org/). The Transcription Factor– Transcription Factor DNA interaction networks were then extracted from the Transcription Factor–gene networks by requiring that both involved nodes of an edge were within the 119 transcription factors studied in Gerstein *et al.*’s work. We studied the hierarchies of these two Transcription Factor–Transcription Factor DNA interaction networks and our Splicing Factor– Splicing Factor mRNA interaction networks with a method similar to that presented in Gerstein *et al.*’s work. In detail, the hierarchy (h) of each node was calculated as h = (Out – In)/(Out + In), where *Out* represents the out-degree and *In* represents the in-degree. A high value means the node is in an “executive” position, while a low level represents the node under regulation. The networks were plotted in three layers: a top “executive” layer (h>1/3), a bottom under-regulation layer (h<-1/3), and a middle layer.

### Assessment of splicing factor binding enrichment to transcripts of splicing factors

Expression data for the HepG2 and K562: ENCODE, whole cell, long poly A, E-GEOD-26284 experiment was downloaded through the EMBL-EBI Expression Atlas (https://www.ebi.ac.uk/gxa/experiments/E-GEOD-26284, accessed on December 12, 2016). Transcripts were sorted by expression, and only transcripts with FPKM (Fragments Per Kilobase of transcript per Million) of 1 or above were included in our analysis. We collected 12,887 genes for K562 and 12,692 for HepG2. Out of the selected 100 splicing regulators, there were 82 splicing factor genes expressed in the HepG2 cell and 84 in the K562 cells with FPKM over 1.

To find the fraction of splicing factors bound by a given splicing regulator, we calculated a ratio of the number of splicing factors a protein binds over the number of expressed splicing factors in a given cell line (K562 or HepG2). We compared this ratio to the number of genes a splicing factor binds over all expressed genes in a given cell line (Figure 5 and Table S3).

## Data availability

eCLIP data sets can be accessed via the ENCODE portal at www.encodeproject.org. Other CLIP data sets can be accessed from the GEO database, with references shown in Table S2.

## Author Contribution

A.D. and S.E.B. conceived of the presented idea. A.D. visualized networks. Z.H. performed the transcription factor and splicing factor hierarchy analyses. A.D. and Z.H. wrote the manuscript with input from J.P.B.L., C.E.F., and S.E.B. Due to an auto accident, S.E.B. has not read the final version of the manuscript.

## Acknowledgements

This work was supported by NIH F31 GM108462 fellowship to A.D, start-up funds to S.E.B, NIH grants (R01 GM071655, U01 EB023686, and U41 HG0077346), and TATA Consultancy Services. We thank Arun Desai for CLIP data download and sorting. We thank Aditya Ramkrishna Rao and his colleagues for help with PubMed search for alternative splicing factor list. We thank Constantina Bakolitsa for reading and commenting on the manuscript.

---

**Figure S1.**
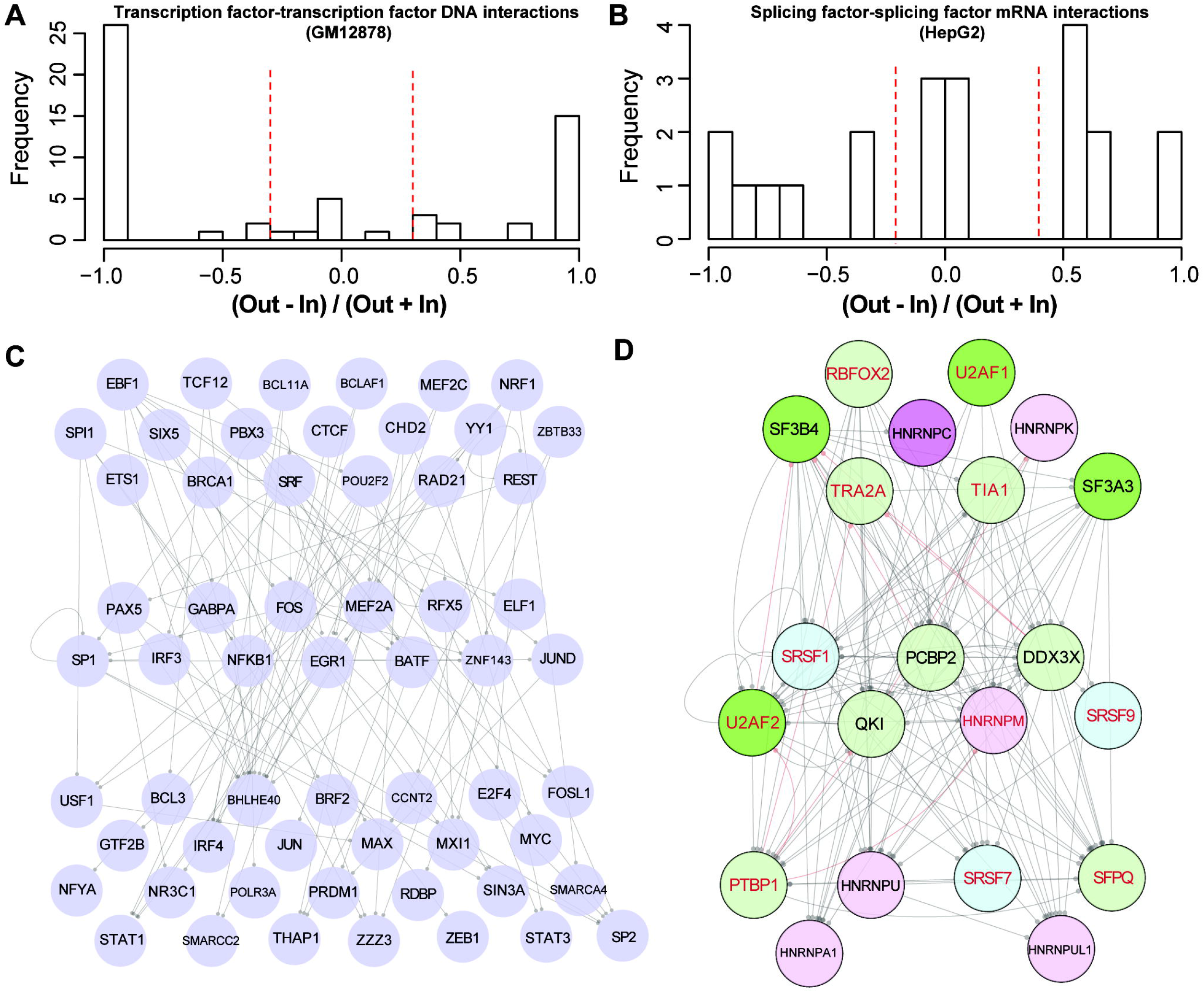
Hierarchies of transcription factor-transcription factor DNA interaction network in GM12878 cell line and splicing factor–splicing factor mRNA interaction network in the HepG2 cell line. **A–B.** The distributions of node hierarchies defined as h = (Out – In)/(Out + In), where “Out” represents the out-degree and “In” represents the indegree, in the transcription factor–transcription factor DNA interaction network (**A**) and splicing factor–splicing factor mRNA interaction network (**B**). **C–D**. The transcription factor– transcription factor DNA interaction network (**C**) and splicing factor–splicing factor mRNA interaction network (**D**) shown in a hierarchical manner. Nodes with h >1/3 and h < −1/3 are on the top and bottom level respectively, with other nodes located in the middle. The grey lines show top-down interactions while red lines show bottom-up interactions. The colors are the same as in Figure 2.

## Supplementary Tables can be found in supplementary excel files

Table S1. Summary of SF-RNA interactions revealed from eCLIP studies

Table S2. Published SF-RNA interactions from CLIP and related studies

Table S3. Summary of splicing factor binding in K562 and HepG2 cell lines

